# A TLR4 ligand-based adjuvant for promoting the immunogenicity of Typhoid subunit vaccines

**DOI:** 10.1101/2024.03.22.581918

**Authors:** Kishore R. Alugupalli

## Abstract

None of the typhoid Vi Polysaccharide (ViPS) subunit vaccines incorporate adjuvants, and the immunogenicity of ViPS vaccines (e.g. Typbar TCV^®^ and Typhim Vi^®^) is in part due to associated TLR4 ligands such as endotoxin present in these vaccines. Since endotoxin content in vaccines is variable and kept very low due to inherent toxicity, I hypothesized that incorporating a defined amount of a non-toxic TLR4-ligand such as monophosphoryl lipid A in ViPS vaccines would improve their immunogenicity. To test this hypothesis, I developed an monophosphoryl lipid A-based adjuvant formulation named Turbo. Admixing Turbo with Typbar TCV^®^ (ViPS-conjugated to tetanus toxoid) increased the levels of anti-ViPS IgM, IgG1, IgG2b, IgG2a/c and IgG3 in inbred and outbred mice. In infant mice, a single immunization with Turbo adjuvanted Typbar TCV^®^, resulted in a significantly increased and durable IgG response, and improved the control of bacterial burden compared to mice immunized without Turbo. Similarly, when adjuvanted with Turbo, the antibody response and control of bacteremia were also improved in mice immunized with Typhim Vi^®^, an unconjugated vaccine. The immunogenicity of unconjugated ViPS is inefficient in young mice and is lost in adult mice when immunostimulatory ligands in ViPS are removed. Nevertheless, when adjuvanted with Turbo, poorly immunogenic ViPS induced a robust IgG response in young and adult mice, and this was observed even under antigen-limiting conditions. These data suggest that incorporation of Turbo as an adjuvant will make typhoid vaccines more immunogenic regardless of their intrinsic immunogenicity or conjugation status and maximize the efficacy across all ages.

## INTRODUCTION

*Salmonella enterica* serovar Typhi (*S.* Typhi) is the causative agent of typhoid fever in humans. There are an estimated 12.5 to 16.3 million cases of typhoid every year with 140,000 deaths. (1). The rapid emergence of multiple drug-resistant strains of *S.* Typhi now complicates the treatment of typhoid (2). Typhoid is a vaccine-preventable disease and vaccination of high-risk populations such as infants and young children is considered the most promising strategy for control (3). Vi polysaccharide (ViPS) of *S.* Typhi is a target for immune responses and anti-ViPS antibodies correlate with protection against typhoid fever (3–5). Two types of ViPS subunit vaccines are currently available (5). Plain ViPS or unconjugated ViPS e.g., Typhim Vi^®^, an US FDA approved vaccine induces T cell-independent B cell response. Its efficacy is ∼55% in older children and adults and the immunity conferred is short-lived (5–7). Importantly, plain ViPS vaccines do not induce antibody responses in children under two years of age. The other type of subunit vaccine is referred to as typhoid conjugate vaccine (TCV), where ViPS is conjugated to protein which induces T cell-dependent B cell responses across all ages. The TCVs utilize a variety of carrier proteins e.g. recombinant Exoprotein A from *P. aeruginosa* (8), CRM197, a non-toxic mutant of diphtheria toxin (9), tetanus toxoid (10), or diphtheria toxoid (11) chemically coupled to ViPS.

None of the typhoid Vi Polysaccharide (ViPS) subunit vaccines incorporate adjuvants, yet these vaccines are immunogenic in both mice and humans. The ViPS antigen is isolated from *Salmonella enterica* serovar Typhi (S. Typhi), a Gram-negative bacterium. Therefore, the ViPS preparation used for making the typhoid subunit vaccines contains trace amounts of bacterial lipopolysaccharide (LPS; also known as endotoxin) a potent activator of Toll-Like Receptor (TLR) 4 signaling which induces co-stimulatory signals. Indeed, it has recently been shown that TLR4 ligands present in Typhoid VIPS subunit vaccines contribute to immunogenicity (12).

Typbar TCV^®^ (ViPS conjugated to tetanus toxoid), the first WHO pre-qualified vaccine shown to be very safe, induces efficient antibody responses in infants and young children with ∼80% efficacy in clinical trials conducted in Malawi, Bangladesh, and Nepal (13–15). The TCV ViPS-CRM197 when tested in a multinational clinical trial in the Philippines, Pakistan and India did not yield an appreciable increase in antibody titers even after two boosters (9, 16). A reason for this hyporesponsiveness to ViPS-CRM197 was in part suggested to be due to the length of the ViPS used for making this TCV (17). Since associated TLR4 ligands present in typhoid vaccines play an important role in the immunogenicity of typhoid subunit vaccines as well as VIPS preparations (12), it is possible that the reduced immunogenicity of ViPS-CRM197 (9) could also be due to a lack of sufficient levels of endogenous TLR4 ligands in the ViPS preparation used for making this vaccine.

Activation of the adaptive immune system requires the engagement of co-stimulatory signaling pathways in addition to B and T cell antigen receptor signaling (18). Adjuvants play a central role by providing co-stimulatory signaling pathways for the immunogenicity of antigens in vaccine formulations. Stimulation of Toll-like receptors (TLRs) triggers the induction of costimulatory molecules, amplifies B cell activation, promotes dendritic cell maturation, and increases antigen-presentation to T cells (19). TLR ligands help direct adaptive immune responses to antigens derived from microbial pathogens, and many vaccines incorporate TLR ligands as adjuvants to augment antigen-specific responses (20). Since TLR4 ligands in typhoid vaccines contribute to immunogenicity (12), in the present study I tested whether incorporating a defined amount of a non-toxic TLR4-ligand, monophosphoryl lipid A-based adjuvant named Turbo in typhoid vaccines would make those vaccines more immunogenic regardless of the difference in the amounts of endotoxin present, the intrinsic immunogenicity of ViPS, or the conjugation of ViPS to a carrier protein.

## MATERIALS AND METHODS

### Mice

The Thomas Jefferson University Institutional Animal Care and Use Committee has approved these studies. Mice were housed in micro-isolator cages with free access to food and water and were maintained in a specific pathogen-free facility. C57BL/6J; (stock no. 000664), 129S1/SvImJ (stock no. 002448), A/J (stock no. 000646), and C3H/HeOuJ (stock no. 000635) were purchased from The Jackson Laboratory (Bar Harbor, ME). CD-1 [Crl:CD1(ICR); Strain code 022] outbred mice were purchased from Charles River Laboratories (Wilmington, MA). Age-matched mice of both sexes were used for all experiments.

### Adjuvant, Antigens, and Immunization

The adjuvant named Turbo was prepared by mixing 1 mg phosphorylated hexaacyldisaccharide (PHAD^®^), a synthetic monophosphoryl Lipid A (MPLA) and 2 mg of 1,2-dipalmitoyl-sn-glycero-3-phosphocholine (Avanti Polar Lipids, Alabaster, AL) in chloroform. Following chloroform evaporation, the contents were suspended in 1% polyethylene glycol sorbitan monooleate (Tween^®^ 80; Sigma-Aldrich) to a concentration of 500 μg/ml of MPLA and homogenized by sonication. The homogenate was mixed 50 times using two syringes connected via a 25G needle and passed 5 times through a polyethersulfone membrane filter with pore size 0.22 μm (Millipore) and the filter-sterilized adjuvant was stored at 4°C. Nanoparticle tracking analysis using NanoSight NS300 instrumentation (Malvern, Instruments Ltd Worcestershire) revealed that the size distribution and concentration of liposomes in the adjuvant formulation were 130 ± 40 nm and 4×10^10^/ml, respectively. The physical characteristics and adjuvant activity was stable for at least one year at 4°C.

ViPS (lot 5, PDMI 158299) was obtained from the U.S. Food and Drug Administration, Silver Spring, MD) (21). A portion of this ViPS preparation was subjected to phenol-extraction to eliminate TLR4 ligands/immunostimulatory compounds as described previously (12). Typhim Vi^®^ (Sanofi Pasteur) an FDA-approved unconjugated typhoid vaccine was purchased, while Typbar TCV^®^ was obtained from Bharat Biotech India Limited, Hyderabad, India.

Mice (3 weeks of age and above) were immunized i.m. with 2.5 μg of ViPS (i.e. Typbar TCV^®^, Typhim Vi^®^, ViPS preparation (US FDA, lot 5, PDMI 158299) without or with phenol extraction) admixed with or without Turbo adjuvant (containing 5 μg MPLA) in 50 μl volume in the thigh region of the hind limb. Infant mice (9-day old) were immunized s.c. with 1.25 μg of ViPS admixed with or without Turbo adjuvant (containing 2.5 μg MPLA) in 25 μl volume in the scruff area between the neck and shoulder. Blood samples were obtained 0, 7-, 14-, 21-28-days (or as indicated) following immunization and stored at −20°C.

### ELISA

ViPS-specific IgM, IgG, IgG1, IgG2b, IgG2a, IgG2c and IgG3 were measured by coating 96-well microtiter plates (Nunc MaxiSorp^TM^; Invitrogen, Carlsbad, CA) with 2 µg/ml of ViPS purified from *S.* Typhi clinical isolate C652464 (22) in DPBS overnight at room temperature. All plates were washed and blocked with 1% Bovine serum albumin in PBS pH 7.2 (blocking buffer) for two hours at room temperature. Blood from ViPS or Typhim Vi^®^ and Typbar TCV^®^ immunized mice was diluted to 1:25 and 1:200, respectively, for IgM and IgG detection. These dilutions were based on a linear range of detection. ViPS-specific mouse IgM, IgG, IgG1, IgG2b, IgG2c, and IgG3 levels were interpreted as ng/μl “equivalents” using normal mouse serum standards (Bethyl Laboratories, Montgomery, TX), mouse isotype specific capture antibodies, and HRPO-conjugated anti-mouse IgM, IgG, IgG1, IgG2b, IgG2c, and IgG3 as described previously (12, 21).

### Infections

To test the relative protection conferred by immunization with Typbar TCV^®^ or Typhim Vi^®^ admixed with and without Turbo adjuvant, mice were infected with a chimeric strain of *S.* Typhimurium (strain RC60) that expresses the *S*. Typhi genes necessary for ViPS synthesis, export and regulation as in *S*. Typhi (23). This protection model was previously validated in several mouse systems (21, 24–26). In brief, Strain RC60 was grown to an OD^600^ of ∼1.0 in Luria-Bertani broth containing 10 mM NaCl. The expression of ViPS was assessed by slide agglutination test using a commercial Vi monoclonal antibody reagent (Statens Serum Institute diagnostica A/S, Denmark; Lot 188L-8). Bacteria were washed twice in DPBS and 100 μl of DPBS containing ∼3×10^4^ CFUs was injected i.p. Three days post-infection liver and spleen were collected, and the tissues were processed using a Minilys tissue homogenizer (Bertin Technologies, Montigny-le-Bretonneux, France). Blood was collected into anti-coagulant and bacterial burden in the blood and tissue homogenates was measured by counting CFUs on LB agar plates.

### Serum bactericidal assay (SBA)

SBA was performed as previously described (27). In brief, log-phase cultures (OD_600_ of 0.5 at 37°C) of *S.* Typhi strain Ty2 were prepared in LB broth with 10 mM NaCl. Bacterial cells were washed in DPBS, and the bacterial cell density was adjusted to 2.5 – 5.0 x 10^4^ colony forming units (CFU) per ml in DPBS. The expression of ViPS was assessed by slide agglutination test using a commercial Vi monoclonal antibody reagent (Statens Serum Institute diagnostica A/S, Denmark; Lot 188L-8). Serum samples were heat-inactivated by incubating at 56°C for 30 minutes prior to use in the assay. Ten microliters of *S*. Typhi strain Ty2 in DPBS (250-500 CFU) were added to each well of a round-bottom polypropylene 96-well plate containing 50 µl of heat-inactivated serum in serial dilutions, 12.5 µl baby rabbit complement (Pel-Freeze, Rogers, AR), and 27.5 µl DPBS. Triplicate samples of each dilution were incubated for 120 minutes at 37°C with gentle rocking and 10 µl of this mixture were plated on LB agar plates for counting CFU. Serum bactericidal antibody titers are defined as the reciprocal of the highest dilution that produced 50% killing in relation to control wells containing complement, but no mouse serum. Naïve mouse serum served as a negative control and serum from either mice immunized with heat-killed *E. coli* strain W3110 expressing pDC5 plasmid, which contains the genes necessary for the synthesis and export of ViPS (28), or *S*. Typhi Anti-Vi human IgG standard (Lot R1, 2011; U.S. Food and Drug Administration, Silver Spring, MD 20993) as two independent positive controls.

### Statistical analysis

Data presented throughout depict pooled data from at least two independent experiments. Statistics were performed using the Prism 10 software program (GraphPad Software, Inc., La Jolla, CA) and the statistical tests are indicated in the figure legends.

## RESULTS

### Turbo adjuvanted Typbar TCV^®^ induces an enhanced anti-ViPS antibody response across all ages

Although Typbar TCV^®^ contains TLR4 ligands or low levels of endotoxin (12), when admixed with Turbo significantly enhanced anti-ViPS IgM, IgG1, IgG2b, IgG2a/c, and IgG3 responses were observed in male and female C57BL6/J mice (Fig. 1). Recently, the immunogenicity of meningococcal conjugate vaccine has been shown to vary among inbred and outbred strains (29). To test the rigor of the immunogenicity promoted by Turbo as an adjuvant we have used outbred strain CD-1 and inbred strains such as C3H and A/J, which can also serve as a model to test vaccine-mediated protection against Salmonella *in vivo* (30). We found that Turbo adjuvanticity occurs in all mouse strains tested, but is strikingly more pronounced in outbred CD-1 strain (Fig. 2).

**Fig. 1.**
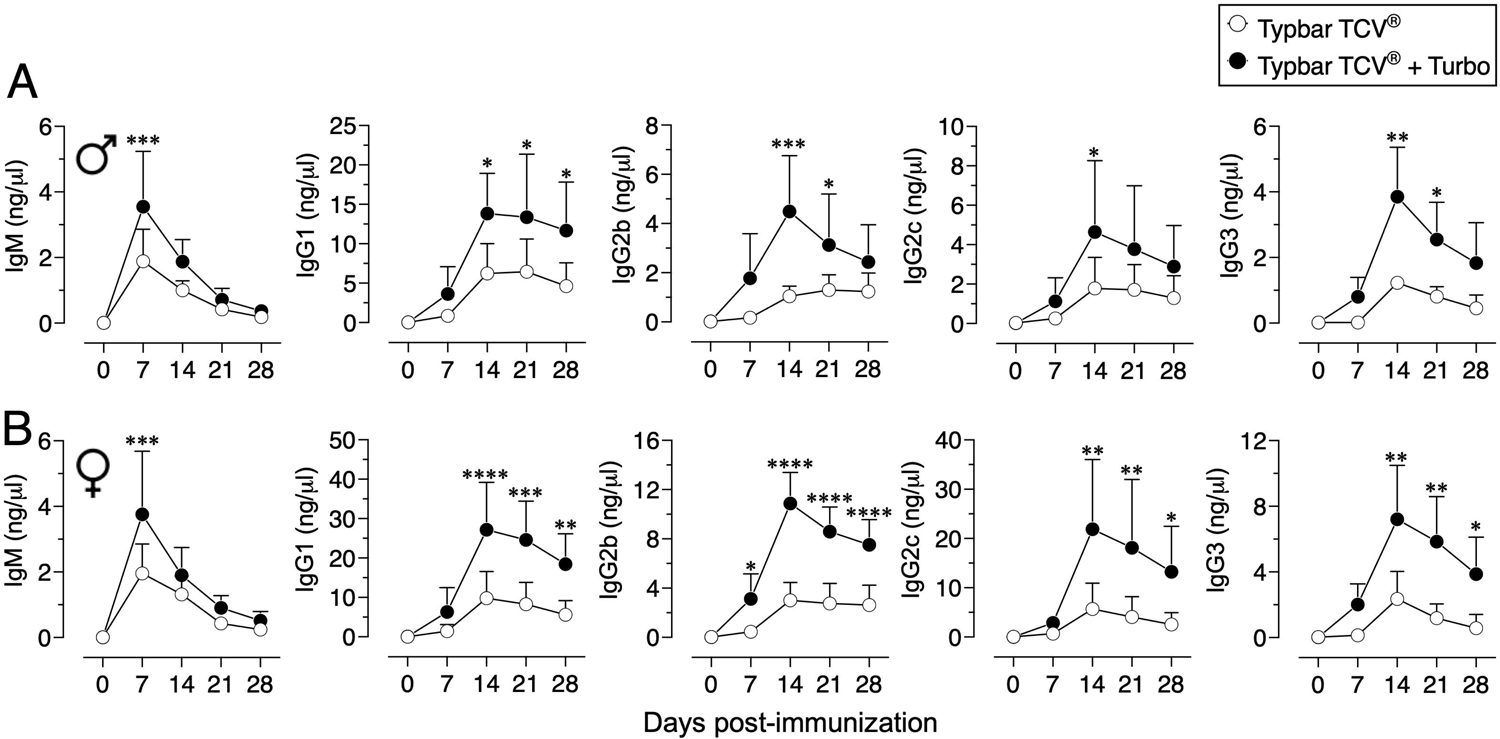
Immunogenicity of Typbar TCV^®^ is enhanced by Turbo. C57BL/6J male (n=6) and female (n=7) 10-12 weeks of age were immunized i.m. with 50 μl of Typbar TCV^®^ vaccines containing 2.5 μg of ViPS admixed with or without Turbo (5μg of MPLA), and ViPS-specific IgM, IgG1, IgG2b, IgG2c, and IgG3 levels were measure by ELISA. Statistics done using 2-way ANOVA Sidak’s multiple comparisons test, and statistically significant differences were indicated as **** = p<0.0001; *** = p<0.001; ** = p<0.001; * = p<0.05.

**Fig. 2.**
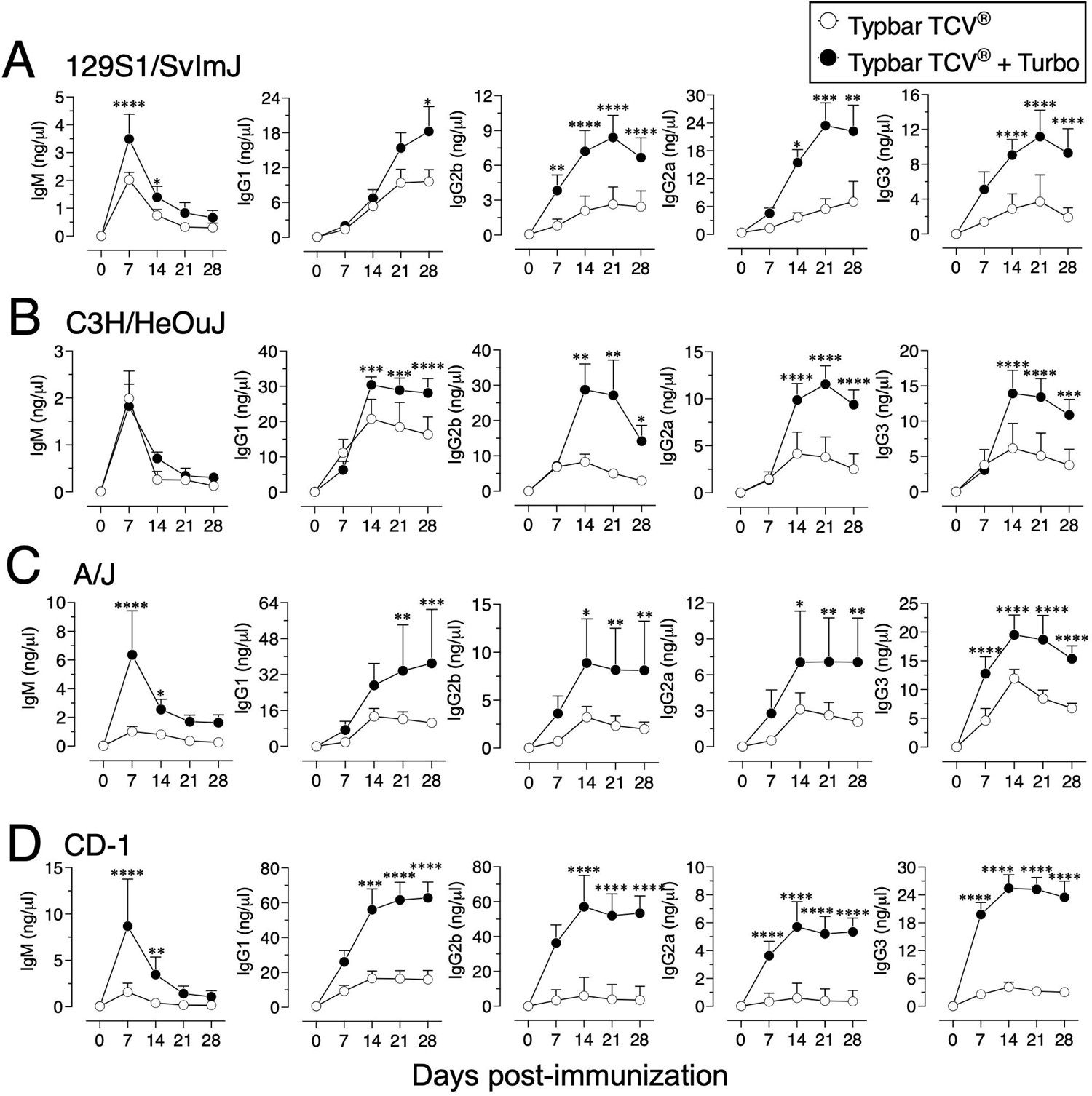
Immunogenicity of Typbar TCV^®^ is enhanced by Turbo in inbred and outbred mice. Inbred mouse strains 129S1/SvImJ, C3H/HeOuJ, A/J or an outbred strain CD-1 of both sexes (n= 6-8) 10-12 weeks age were immunized i.m. with 50 μl of Typbar TCV^®^ vaccines containing 2.5 μg of ViPS admixed or without Turbo (5μg of MPLA), and ViPS-specific IgM, IgG1, IgG2b, IgG2a, and IgG3 levels were measure by ELISA. Statistics done using 2-way ANOVA Sidak’s multiple comparisons test, and statistically significant differences were indicated as **** = p<0.0001; *** = p<0.001; ** = p<0.001; * = p<0.05.

Aged individuals do not respond efficiently to vaccines due to immune senescence (31), and the anti-ViPS response is lower in 82-week-old mice compared to that in 16-week-old adult mice (Fig 3A vs B). However, when adjuvanted with Turbo, aged mice showed a significant improvement in anti-ViPS IgM and IgG responses to Typbar TCV^®^ (Fig. 3A vs B). The lack of an efficient antibody response early in life is due to an incomplete development of the immune system (32). Nevertheless, when Typbar TCV^®^ is adjuvanted with Turbo, infant mice produced a rapid IgM response and an order of magnitude higher IgG response that is sustained for at least 120 days post-immunization compared to mice that were immunized with Typbar TCV^®^ alone (Fig. 3C). Since S. Typhi does not infect commonly used mouse strains (33), and we have used S. Typhimurium strain RC60 expressing ViPS antigen to test ViPS-mediated protection *in vivo,* as described previously (21, 24–26). We found that infant mice vaccinated with Turbo-adjuvanted Typbar TCV^®^, when challenged 120 days-post immunization with S. Typhimurium strain RC60, exhibited more significant reduction in bacterial load in the blood, liver, and spleen, compared to mice immunized without Turbo (Fig. 3D).

**Fig. 3.**
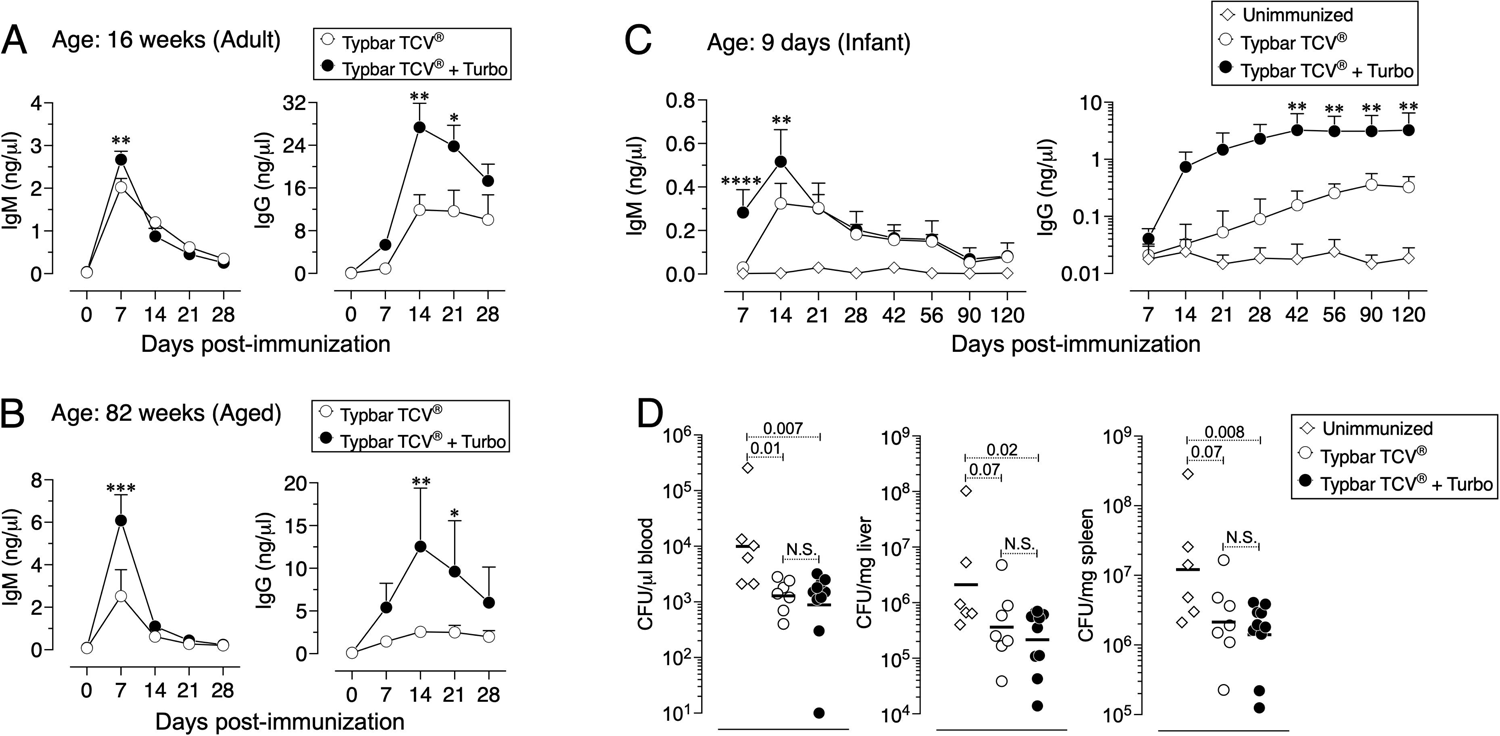
Adjuvanticity of Turbo occurs across all ages of mice. C57BL/6J mice of 16 (A) or 82 (B) weeks of age (adult and aged) (n=8) were immunized i.m. with 50 μl of Typbar TCV^®^ vaccines containing 2.5 μg of ViPS admixed with or without Turbo (5μg of MPLA). (C) Infant mice (9 day old) were immunized s.c. with 1.25 μg of ViPS admixed with or without Turbo adjuvant (containing 2.5 μg MPLA) in 25 μl volume in the scruff area between neck and shoulder. ViPS-specific IgM and IgG levels were measured by ELISA and statistics were done using 2-way ANOVA Sidak’s multiple comparisons test and statistically significant differences were indicated as **** = p<0.0001; *** = p<0.001; ** = p<0.001; * = p<0.05. (D) After 120 days post-immunization of mice in panel C were infected with with 3 x 10^4^ CFU of ViPS expressing *S*. Typhimurium strain RC60 intraperitoneally. Three days post-infection mice were sacrificed and bacterial burden in the blood, liver and spleen was determined by plating serial 10-fold dilutions of blood or tissue homogenates followed by colony counting. Each dot represents an individual mouse, and the black bar represents mean. Statistical differences were determined by Mann-Whitney U test.

### The unconjugated ViPS vaccine, Typhim Vi^®^ adjuvanted with Turbo induces enhanced antibody responses

Compared to conjugated vaccines such as Typbar TCV^®^, the antibody responses to Typhim Vi^®^, is an order of magnitude lower (12). Despite the presence of TLR4 ligands in Typhim Vi^®^ (12), when this vaccine was adjuvanted with Turbo, all IgG isotypes were enhanced (Fig. 4A). Moreover, seroconversion with the adjuvanted Typhim Vi^®^ was 100%, unlike the unadjuvanted Typhim Vi^®^, and IgG responses were sustained for at least 90 days (Fig. 4B), and the control of bacteremia was significantly improved (Fig. 4C).

**Fig. 4.**
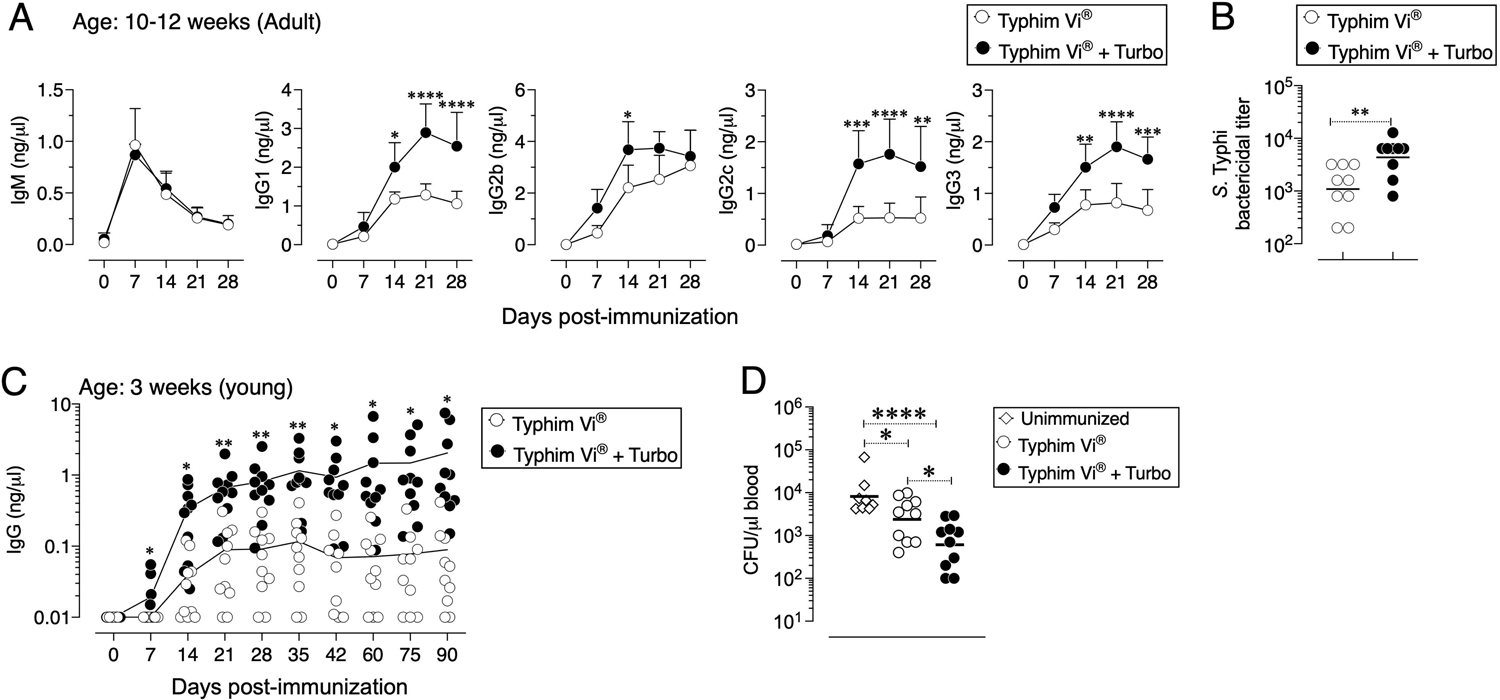
Immunogenicity of unconjugated vaccine Typhim Vi^®^ is enhanced by Turbo. (A) C57BL/6J mice of 10-12-weeks (n=9) and (C) 3-weeks (n=10) of age were immunized i.m. with 50 μl of Typhim Vi^®^ containing 2.5 μg of ViPS admixed with or without Turbo (5μg of MPLA), and ViPS-specific IgM, IgG, IgG1, IgG2b, IgG2c, and IgG3 levels were measure by ELISA. Statistics done using 2-way ANOVA Sidak’s multiple comparisons test, and statistically significant differences were indicated as **** = p<0.0001; *** = p<0.001; ** = p<0.001; * = p<0.05. (C) Serum bactericidal antibody titers against *S.* Typhi strain Ty2 were determined using serum obtained from 28-days post-immunization. Each dot represents an individual mouse, and the bar represents geometric mean. Statistical differences were determined by Mann-Whitney U test. (D) After 90 days post-immunization of mice in panel C were infected with with 3 x 10^4^ CFU of ViPS expressing *S*. Typhimurium strain RC60 intraperitoneally. Three days post-infection mice were sacrificed and bacterial burden in the blood was determined by plating serial 10-fold dilutions of blood followed by colony counting. Each dot represents an individual mouse, and the black bar represents mean. Statistical differences were determined by Mann-Whitney U test.

### ViPS and poorly immunogenic ViPS preparations when adjuvanted with Turbo induce a dramatically improved IgM and IgG responses in young and adult mice

Unconjugated polysaccharide vaccines such as Typhim Vi^®^ do not induce an efficient response in young (3-week-old) compared to adult mice due to a restricted V_H_ gene usage early in life and a paucity of antigen-specific B cell precursors (26). To test whether admixing Turbo with ViPS (US FDA, lot 5, PDMI 158299) overcomes the poor antibody responses in the young, 3-week-old C57BL/6J mice were immunized. We found that Turbo-adjuvanted ViPS dramatically improved the IgG response in young mice (Fig. 5A). We showed that this ViPS preparation contains TLR4 ligands that play an important role in the immunogenicity of this ViPS (12), and that the elimination of these immunostimulatory components by phenol extraction results in a dramatic loss of immunogenicity (12). Remarkably, when phenol extracted ViPS preparation was admixed with Turbo, immunogenicity was restored in young mice (Fig. 5B). Furthermore, we found that native ViPS and phenol extracted ViPS preparation when adjuvanted with Turbo induced efficient IgM and all four IgG isotypes in adult inbred and outbred strains of mice compared to mice immunized without Turbo (Fig 6). These data suggest that Turbo can serve as an adjuvant for unconjugated ViPS vaccines in young and adult mice.

**Fig. 5.**
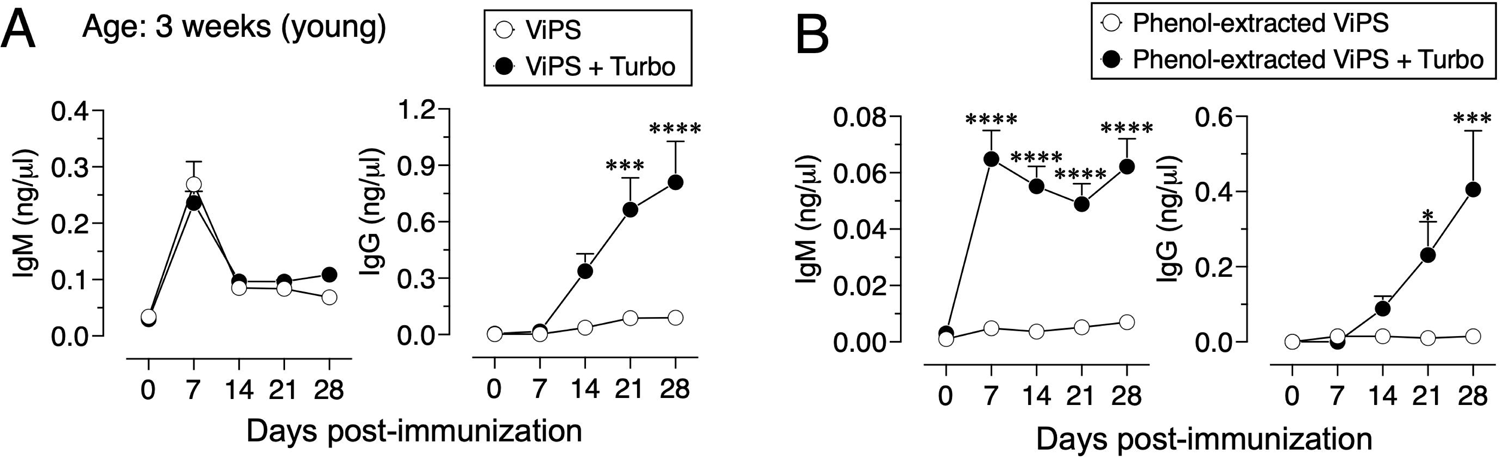
Admixing Turbo promoted poorly immunogenic ViPS in young mice. C57BL/6J mice of 3 weeks of age (n=6) were immunized i.m. with 50 μl 2.5 μg of (A) ViPS (lot 5 PDML158299 from US FDA), or (B) same ViPS stock subjected to phenol extraction to eliminate TLR4 ligand, were admixed with or without Turbo (5μg of MPLA), and ViPS-specific IgM and IgG levels were measure by ELISA. Statistics done using 2-way ANOVA Sidak’s multiple comparisons test, and statistically significant differences were indicated as **** = p<0.0001; *** = p<0.001; * = p<0.05.

**Fig. 6.**
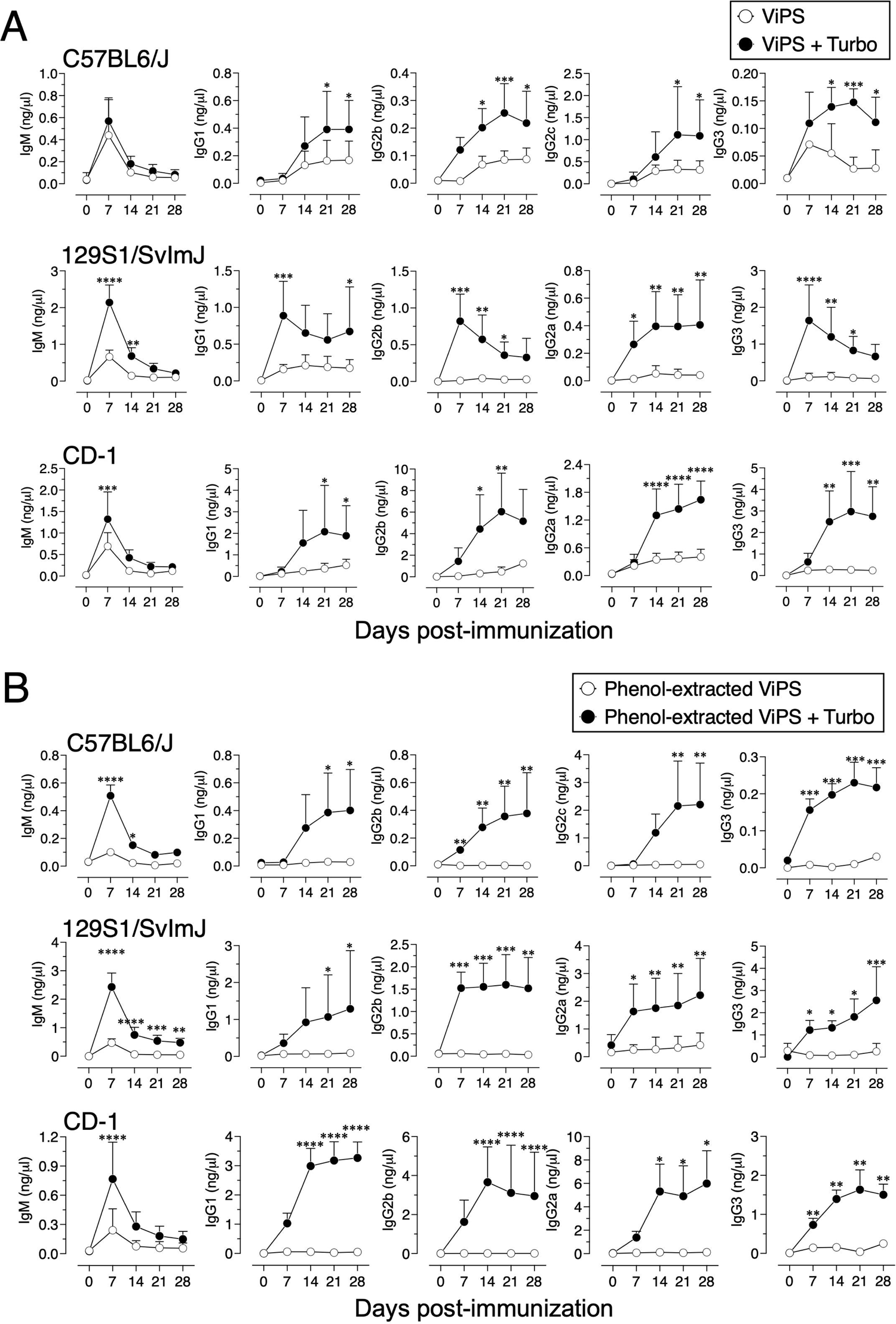
Admixing Turbo promoted immunogenicity of ViPS in adult inbred and outbred mouse strains. C57BL/6J, 129S1/SvImJ and CD-1 mice of 9-12 weeks of age representing both sexes (n=6-8) were immunized i.m. with 50 μl 2.5 μg of (A) ViPS (lot 5 PDML158299 from US FDA), or (B) phenol-extracted ViPS were admixed with or without Turbo (5μg of MPLA), and ViPS-specific IgM, IgG1, IgG2b, IgG2a/c, and IgG3 were measure by ELISA. Statistics done using 2-way ANOVA Sidak’s multiple comparisons test, and statistically significant differences were indicated as **** = p<0.0001; *** = p<0.001; ** = p<0.01 * = p<0.05.

### Turbo adjuvanicity occurs under antigen-limiting conditions

A characteristic of adjuvants is their ability to promote antigen-specific responses even at very low concentrations of a given antigen. To test the impact of Turbo adjuvanicity on dose-sparing of the antigen we immunized mice with decreasing amounts of ViPS over a 100-fold range (2.5 to 0.02μg). We found that admixing Turbo promoted immune responses even at a concentration (0.02μg) that failed to induce a detectable anti ViPS-IgG response without Turbo (Fig. 7A). We also found that at a constant concentration of ViPS (2.5μg), Turbo containing 5 μg of MPLA induced the highest IgM and IgG response, suggesting that this amount of MPLA is optimal in mice.

**Fig. 7.**
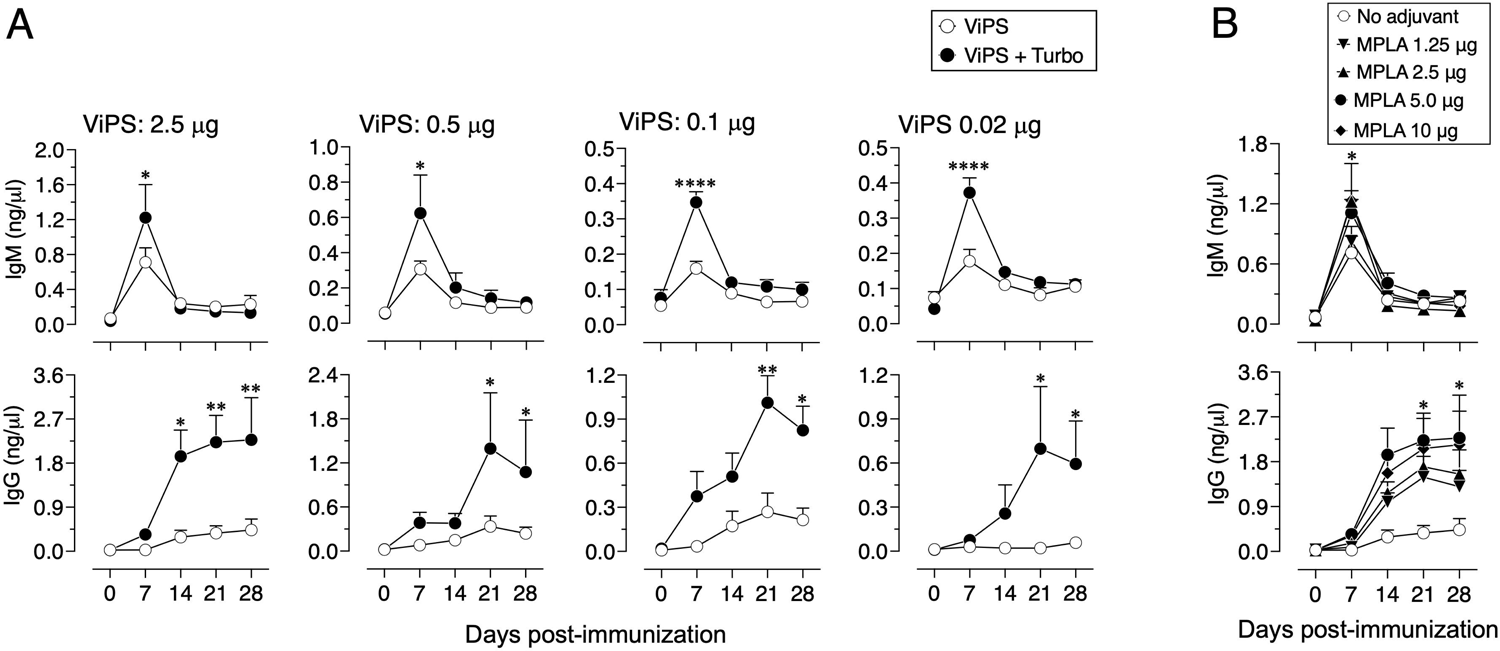
Turbo adjuvanticity under limiting concentrations of ViPS antigen. (A) C57BL6 mice of 10-week-old both sexes (n=3 each) were immunized i.m. with 2.5, 0.5, 0.1, 0.02 μg of ViPS in the presence or absence of Turbo (5mg MPLA). (B) 2.5μg of ViPS in the presence varying concentrations of MPLA in Turbo. ViPS-specific IgM and IgG were measured by ELISA. Statistics done using 2-way ANOVA Sidak’s multiple comparisons test, and statistically significant differences were indicated as **** = p<0.0001; *** = p<0.001; ** = p<0.01 * = p<0.05.

## DISCUSSION

The major advantage of subunit vaccines is their safety. However, one of their major disadvantages is their inability to induce efficient and long-lasting immunity against disease. Therefore, incorporation of adjuvants is required for the immunogenicity of subunit vaccines. Typhim Vi^®^ has an efficacy of appproximately 55% in older children and adults, and the immunity conferred is short-lived. In fact, seropositivity in individuals drops to 46% by two years post-vaccination (5–7). Despite having TLR4 ligand activity in Typhim Vi^®^ (12), in the present study I found that when adjuvanted with Turbo the immune response to Typhim Vi^®^ increases significantly and the seroconversion sustains at 100% for as long as 90 days post-immunization and confers a significant protection (Fig 4). Single immunization with Typbar TCV^®^ induces very efficient antibody responses in mice (12), and the efficacy of Typbar TCV^®^ was shown to be approximately 80% in three independent clinical trails (13–15). Typbar TCV^®^ is the first prequalified TCV by the WHO in 2017. Using Typbar TCV^®^ as a comparator, another TCV, TyphiBEV^®^ demonstrated its equivalence to Typbar TCV^®^ in terms of immunogenicity, reactogenicity, and safety in healthy infants, children, and adults (34).

None of the typhoid subunit vaccines including TyphiBEV^®^, the second prequalified TCV by the WHO in 2020, contain adjuvants. Recently I have shown that antibody responses to Typbar TCV^®^ and Typhim Vi^®^ is dependent on TLR4 ligands present in these vaccines (12). The ViPS used for TyphiBEV^®^ is isolated from *Citrobacter freundii* strain WR7011 (35), and is structurally identical to the ViPS from *S.* Typhi (36). In the present study I have used the same ViPS preparation (US FDA, lot 5, PDMI 158299) that was shown to contain TLR4 ligands (12) to test the adjuvanticity of Turbo (Figs 5, 6 & 7). Therefore, TyphiBEV^®^ is expected to contain TLR4 ligands and its immunogenicity is also expected to be dependent on these ligands as seen with Typbar TCV^®^ (12). In the present study I show that mixing of Turbo with Typbar TCV^®^ or Typhim Vi^®^ significantly improves antigen-specific IgG response across all ages. The lack of an efficient antibody response, particularly to bacterial polysaccharide antigens, in human infants and infant mice is due to an incomplete B cell development and restricted BCR repertoire (37). Consistent with this we have previously shown that in IL-7 dependent B lymphopoiesis and distal V_H_ gene usage are required for the anti-ViPS antibody response in mice (26). Studies in mice have shown that the impairment in immune responses in aged animals is due to several immunological parameters including a reduced B cell antigen receptor (BCR) repertoire (38), and decreased numbers of antigen-presenting cells and their interaction with T cells (39, 40). An improved response in infant and aged mice when adjuvanted with Turbo (Fig. 3) suggests that the admixing Turbo increases the antigen-specific specific B cell precursor frequency. Therefore, incorporation of Turbo as an adjuvant with typhoid subunit vaccines is expected to improve the magnitude and durability of the immune response and efficacy of the typhoid vaccines across all ages.

Many other Gram-negative bacterial polysaccharide vaccines such as those against *Neisseria meningitidis* and *Hemophilus influenzae* type B (HiB) e.g. FDA-approved Menveo^®^, MenQuadfi^®^, ActHIB^®^ and HIBERIX^®^, do not incorporate adjuvants. Although the endotoxin levels in these vaccines are within the acceptable limits allowed by regulatory agencies, the levels are expected to vary from lot to lot as seen with Hib vaccines (41). In fact, I have shown that a variation in TLR4 ligands can account for the differences in the immunogenicity of unadjuvanted vaccines (12) (Fig. 5 & 6). Unlike the immunogenicity seen with TCVs with a single immunization in infants (13–15), the induction of an optimal antibody responses by HiB and meningococcal polysaccharide conjugate vaccines require up to three boosters at ages 4, 6, and 12-15 months. This multiple booster immunization strategy is not only cost-prohibitive but also a challenge for compliance particularly for low- and middle-income countries (LMICs). The development of adjuvanted polysaccharide conjugate vaccines that are more immunogenic, less expensive, and require fewer doses will limit visits to the clinics for periodic boosting would result in greater global accessibility and higher compliance in the broader population.

The inclusion of MPLA as an adjuvant component is approved by the FDA for several viral vaccines e.g. Zoster vaccine (Shingrix^®^), Human papillomavirus vaccine (Cervarix^®^), and Hepatitis B vaccine (Fendrix^®^). In addition to MPLA, these vaccines also contain other adjuvant components such as QS21, a fraction extracted from Quillaja saponaria plant, and Al(OH)3 or AlPO4 commonly referred as Alum (42). While MPLA is considered as an adjuvant for protein antigens, the impact of MPLA-based adjuvants was not thoroughly explored for bacterial polysaccharide vaccines. Recently it was shown that incorporation of a mixture of adjuvants that activate TLR, C-type lectin receptor, and squalene was shown to enhance antibody responses to Pneumovax^®^23, suggesting the utility of adjuvants for unconjugated pneumococcal polysaccharide vaccines in improving their immunogenicity (43). The present study suggests that incorporating Turbo in bacterial polysaccharide subunit vaccines could help reduce the amount of antigen needed, minimize boosters for vaccines such as meningococcal and HiB vaccines, and make all vaccines immunogenic regardless of the antigen isolation process, intrinsic antigenicity, or the carrier protein or method used for polysaccharide conjugation and maximize the immunogenicity and efficacy across all ages.

## ACKNOWLEDGEMENTS

I thank Dr. Krishna Mohan of Bharat Biotech India Limited for providing the Typbar TCV vaccine, and Dr. John Cipollo of US FDA and Dr. Sudeep Kothari of IVI for providing ViPS. I also thank Darren Dougharty for helping in blood sampling of mice, and Dr. Tim Manser for comments and critical reading of this manuscript.

## AUTHOR CONTRIBUTIONS

K.R.A. conceived the hypothesis, designed, and performed the experiments, analyzed the data, and wrote the manuscript.

